# Lysosome enlargement during inhibition of the lipid kinase PIKfyve proceeds through lysosome coalescence

**DOI:** 10.1101/295246

**Authors:** Christopher H. Choy, Golam Saffi, Matthew A. Gray, Callen Wallace, Roya M. Dayam, Zhen-Yi A. Ou, Guy Lenk, Rosa Puertollano, Simon C. Watkins, Roberto J. Botelho

**Author notes:** Equal contribution (Christopher H. Choy, Golam Saffi). To whom correspondence should be sent: Address: Department of Chemistry and Biology, Ryerson University, Toronto, ON, M5B2K3, Canada.

## Abstract

Lysosomes receive and degrade cargo from endocytosis, phagocytosis and autophagy. They also play an important role in sensing and instructing cells on their metabolic state. The lipid kinase PIKfyve generates phosphatidylinositol-3,5-bisphosphate to modulate lysosome function. PIKfyve inhibition leads to impaired degradative capacity, ion dysregulation, abated autophagic flux, and a massive enlargement of lysosomes. Collectively, this leads to various physiological defects including embryonic lethality, neurodegeneration and overt inflammation. While being the most dramatic phenotype, the reasons for lysosome enlargement remain unclear. Here, we examined whether biosynthesis and/or fusion-fission dynamics contribute to swelling. First, we show that PIKfyve inhibition activates TFEB, TFE3 and MITF enhancing lysosome gene expression. However, this did not augment lysosomal protein levels during acute PIKfyve inhibition and deletion of TFEB and/or related proteins did not impair lysosome swelling. Instead, PIKfyve inhibition led to fewer but enlarged lysosomes, suggesting that an imbalance favouring lysosome fusion over fission causes lysosome enlargement. Indeed, conditions that abated fusion curtailed lysosome swelling in PIKfyve-inhibited cells.

**Summary statement:** PIKfyve inhibition causes lysosomes to coalesce, resulting in fewer, enlarged lysosomes. We also show that TFEB-mediated lysosome biosynthesis does not contribute to swelling.

## Introduction

Lysosomes are organelles responsible for the degradation of a large variety of molecules that originate from within and from outside cells. Lysosomes accomplish this by fusing with cargo-containing organelles such as autophagosomes, Golgi-derived vesicles, endosomes and phagosomes (Luzio et al., 2007, 2009; Schwake et al., 2013). To enable their degradative capacity, lysosomes are equipped with a plethora of hydrolytic enzymes that digest proteins, lipids, nucleic acids, and carbohydrates. In addition, the vacuolar H^+^-pump ATPase generates a highly acidic lumen optimal for hydrolytic activity (Mindell, 2012; Schwake et al., 2013; Xiong and Zhu, 2016; Luzio et al., 2007). Nonetheless, the function of lysosomes is not limited to molecular degradation. Among other roles, lysosomes are an important storage organelle, they mediate antigen presentation, they undergo exocytosis to repair membrane damage, and are a genuine signalling platform that senses and informs the cell of its nutrient and stress conditions (Medina et al., 2015; Settembre and Ballabio, 2014; Sancak et al., 2010; Delamarre et al., 2005; Xiong and Zhu, 2016; Reddy et al., 2001; Settembre et al., 2012; Roczniak-Ferguson et al., 2012).

Strikingly, lysosomes are now considered a highly dynamic and heterogeneous organelle. Not only can individual cells host tens to hundreds of lysosomes, lysosomes themselves are highly heterogeneous in their pH (Johnson et al., 2016), subcellular position (Li et al., 2016a; Pu et al., 2015; Bagshaw et al., 2006), and morphology (Swanson et al., 1987; Saric et al., 2016; Vyas et al., 2007). Lastly, lysosomes can adapt to their environment. For example, in macrophages and dendritic cells, lysosomes are transformed from individual puncta into a highly tubular network (Saric et al., 2016; Vyas et al., 2007; Swanson et al., 1987). In addition, nutrient depletion, protein aggregation, and phagocytosis activate a family of transcription factors that can enhance expression of lysosome genes and adjust lysosome activity (Gray et al., 2016; Pastore et al., 2016; Sardiello et al., 2009; Tsunemi et al., 2012; Medina et al., 2011; Settembre et al., 2012; Roczniak-Ferguson et al., 2012). This is best understood for the related transcription factors, TFEB and TFE3, which are recruited to the surface of lysosomes and subject to phosphorylation by various kinases including mTORC1 to modulate their nucleo-cytoplasmic transport (Li et al., 2016b; Martina et al., 2012; Peña-Llopis et al., 2011; Martina et al., 2014, 2016; Roczniak-Ferguson et al., 2012).

Given the above, lysosomes are highly regulated organelles. A master modulator of lysosomes is the lipid kinase PIKfyve, which converts phosphatidylinositol-3-phosphate [PtdIns(3)P] into phosphatidylinositol-3,5-bisphosphate [PtdIns(3,5)P_2_] (Sbrissa et al., 1999). Inactivation of PIKfyve or of its regulators, Vac14/ArPIKfyve and Fig4/Sac3, causes a variety of physiological problems including embryonic lethality, neurodegeneration and immune malfunction (Cai et al., 2013; Chow et al., 2007; Jin et al., 2008; Ho et al., 2012; Mccartney et al., 2014; Sbrissa et al., 2007; Ferguson et al., 2010; Min et al., 2014; Lenk et al., 2016; Ikonomov et al., 2011). At the cellular level, these defects relate to impaired autophagic flux, altered delivery to lysosomes, dysregulation of lysosomal Ca^2+^ transport, and massive enlargement of lysosomes (Ferguson et al., 2009; Dong et al., 2010; de Lartigue et al., 2009; Ferguson et al., 2010; Kim et al., 2014; Ho et al., 2012; Mccartney et al., 2014; Kim et al., 2016). Remarkably, it remains unclear how lysosomes enlarge in cells defective for PIKfyve activity. It has been suggested that PIKfyve, or loss of the yeast ortholog, Fab1, impairs recycling from endosomes, lysosomes or from the yeast vacuole, thus causing an influx of membrane into lysosomes and their enlargement (Rutherford et al., 2006; Bryant et al., 1998; Dove et al., 2004). This recycling process is assumed to occur through vesicular-tubular transport intermediates like those induced by clathrin, adaptor proteins and retromer (Currinn et al., 2016; de Lartigue et al., 2009; Ho et al., 2012; Phelan et al., 2006), but remains speculative. This type of membrane fission is distinct from “kiss-and-run” process, where lysosomes undertake transient homotypic and heterotypic fusion events that are followed by fission to avoid coalescence of lysosomes (Bright et al., 2005). Membrane fission can also occur as organelle splitting, rather than budding, as is the case of mitochondria fission (Chan, 2006).

Recently, PIKfyve impairment was shown to cause nuclear accumulation of TFEB, suggesting that increased lysosomal biogenesis may contribute to lysosome swelling (Gayle et al., 2017; Wang et al., 2015). Here, we examined the role of TFEB and the related TFE3 and MITF in lysosome enlargement and conclude that biosynthesis does not contribute to lysosome enlargement, at least acutely. Instead, we observed that PIKfyve inhibition causes a significant reduction in lysosome number concurrent with lysosome enlargement. We propose that lysosome swelling proceeds through lysosome coalescence, likely caused by reduced “run”/fission events during lysosome “kiss-and-run” and/or full fusion/fission cycles.

## Results

### PIKfyve inhibition activates the transcription factor TFEB independently of mTOR

PIKfyve inhibition causes lysosome enlargement (Ikonomov et al., 2001; Mccartney et al., 2014). Nevertheless, the mechanism by which this happens remains unclear. There is a growing body of literature that links TFEB and related transcription factors to enhanced lysosome and protein expression (Settembre and Ballabio, 2014; Raben and Puertollano, 2016). Thus, we set out to test if TFEB activation might contribute to lysosome swelling. Indeed, PIKfyve inhibition with apilimod, an exquisitely specific inhibitor of PIKfyve (Cai et al., 2013; Gayle et al., 2017), in RAW cells triggered a rapid and robust nuclear translocation of GFP-tagged TFEB and endogenous TFEB (Fig. 1A, B). To complement our pharmacological treatment, we also demonstrated that siRNA-mediated silencing of PIKfyve significantly boosted the number of RAW cells with nuclear TFEB-GFP relative to control-silenced cells (Fig. 1C). In addition, HeLa cells treated with apilimod also exhibited nuclear TFEB relative to resting cells, showing that activation of TFEB in the absence of PIKfyve occurs across distinct cell types (Fig. 1D). Lastly, we showed that GFP-fusion of TFE3 and MITF, which are related to TFEB (Ploper and De Robertis, 2015; Raben and Puertollano, 2016), also translocated to the nucleus upon PIKfyve abrogation in RAW cells (Supplemental Fig. S1A,B). Overall, our data is consistent with recent reports that PIKfyve activity robustly controls nuclear localization of TFEB and related factors (Gayle et al., 2017; Wang et al., 2015).

**Figure 1:**
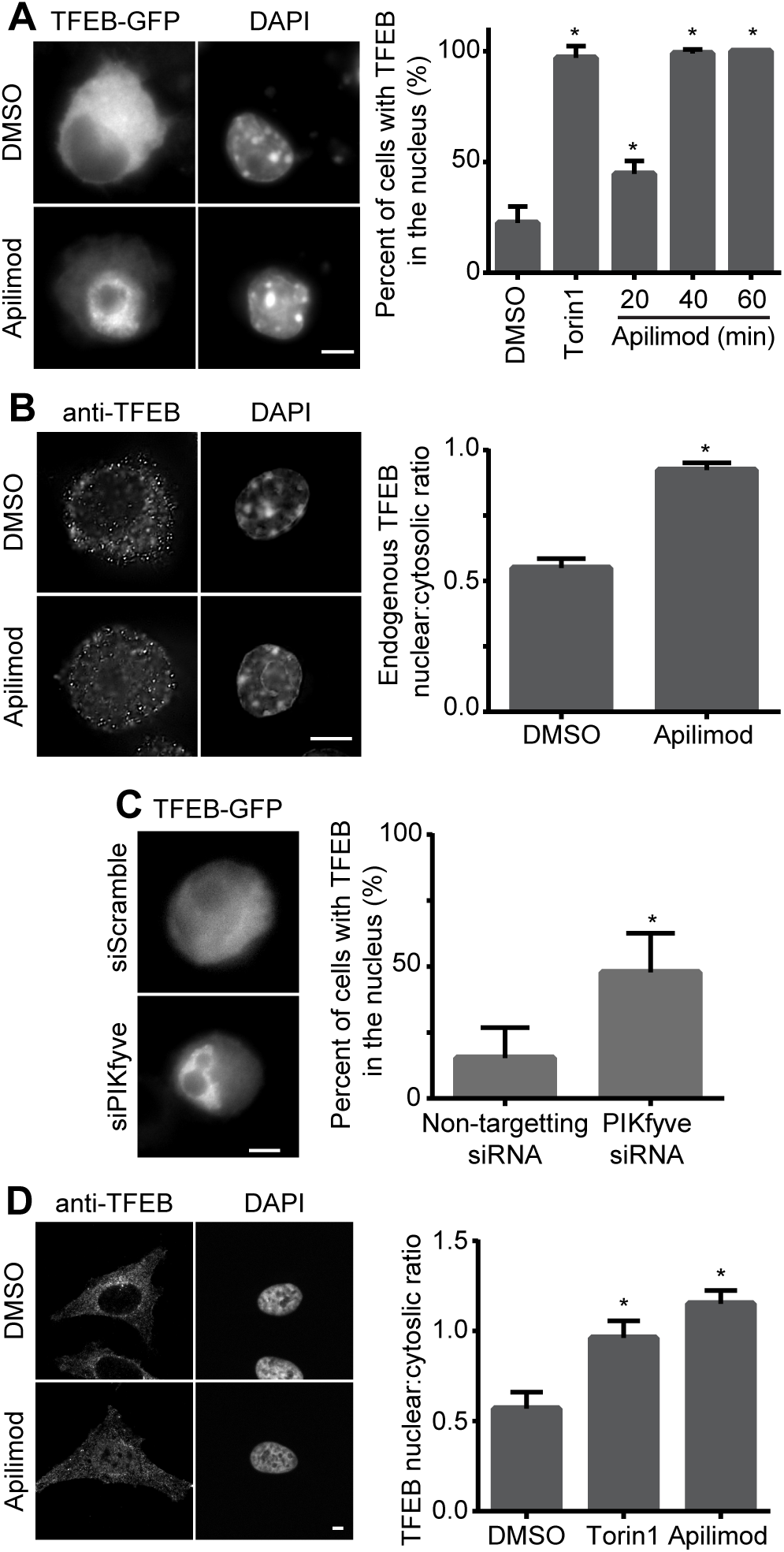
PIKfyve inhibition causes nuclear translocation of TFEB. A, B: RAW cells expressing TFEB-GFP (A) or stained for endogenous TFEB (B) treated with vehicle or 20 nM apilimod for 1 h. C: RAW cells silenced for PIKfyve or mock-silenced and expressing TFEB-GFP. D: HeLa cells stained for endogenous TFEB treated with vehicle or 20 nM apilimod for 1 h. Where shown, DAPI stains the nuclei. Nuclear translocation of TFEB was quantified either as percent of cells containing nuclear GFP-tagged TFEB (A, C) or as the nuclear-to-cytosol fluorescence intensity ratio for endogenous TFEB (B, D). Shown is the mean and standard error of the mean from at least three independent experiments and based on 50-300 cells per condition per experiment. Asterisks (*) indicate statistical difference from respective control conditions (p<0.05) using one-way ANOVA and Tukey’s post-hoc test or with an unpaired Student’s t-test, as appropriate. Scale bar = 5 µm.

TFEB is controlled by several factors including mTOR, which phosphorylates and maintains TFEB in the cytosol (Martina et al., 2012; Roczniak-Ferguson et al., 2012; Settembre et al., 2012). In addition, PIKfyve has been suggested to control mTORC1 in adipocytes (Bridges et al., 2012), though others have noted that mTOR activity is sustained in PIKfyve inhibited cells (Wang et al., 2015; Krishna et al., 2016). To better assess the possibility that PIKfyve controls TFEB by coordinating with mTOR in our cells, we examined the phosphorylation state of S6K, a major mTORC1 target, in cells treated with apilimod. Strikingly, we could not observe a significant difference in the phosphorylation levels of S6K between control and apilimod-treated cells (Fig. 2A). This suggests that mTORC1 retains its activity in PIKfyve-inhibited RAW macrophages. Alternatively, we questioned if mTOR might control TFEB by regulating PIKfyve activity instead. To test this, we exposed cells to torin1, an inhibitor of mTOR. First, we showed that apilimod reduced PtdIns(3,5)P_2_ by >70% with a concomitant increase in its precursor phosphatidylinositol-3-phosphate (Fig. 2B). In comparison, we did not observe a decrease in the levels of both phosphatidylinositol-3-phosphate or PtdIns(3,5)P_2_ in torin1-treated cells relative to control cells; in fact, there was a statistically significant increase in PtdIns(3,5)P2 levels in torin1-treated cells (Fig. 2B).

**Figure 2:**
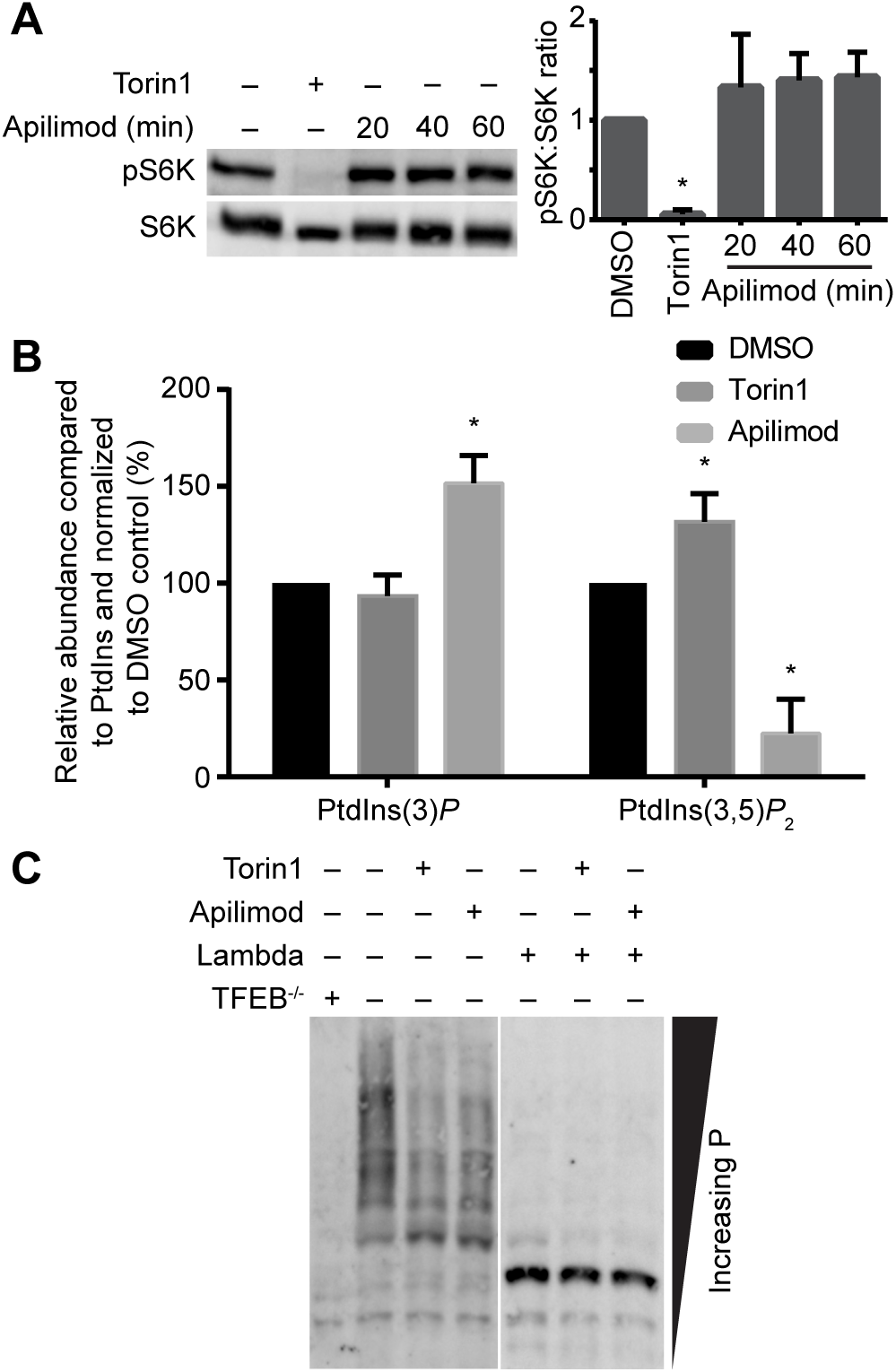
mTOR and PIKfyve function independently. A. Western blot detection of phosphoThr^389^-p70S6K and total p70S6K in PIKfyve-inhibited cells treated as indicated. The ratio of phosphor-S6K-to-total S6K was quantified from three independent blots. Shown is the mean ratio and standard deviation normalized to DMSO-treated cells. B. Quantification of PtdIns(3)P and PtdIns(3,5)P_2_ levels using ^3^H-*myo*-inositol incorporation and HPLC-coupled flow scintillation. Results are shown as the mean and standard deviation from three independent experiments, normalized to their PtdInsP levels and to the respective DMSO-treated controls. For A and B, asterisks (*) indicate statistical difference from respective control conditions (p<0.05) using multiple Student’s t-test with the Bonferroni correction. C. Western blot against TFEB from whole cell lysates resolved by a Phos-tag gel. Shown in a representative blot from three independent experiments showing differential migration of TFEB in lysates from control relative to torin1- and apilimod-treated cells. Anti-TFEB antibody specificity was confirmed by the loss of most bands in *tfeb*^−/-^ RAW whole cell lysates. In addition, most bands represent phosphorylated isoforms of TFEB since pre-treatment of lysates with lambda protein phosphatase led to a single fast-running band.

Since TFEB is regulated by phosphorylation at various sites (Dephoure et al., 2008; Mayya et al., 2009), we decided to assess whether PIKfyve activity might govern TFEB through phosphorylation. To achieve this, we examined changes in TFEB phosphorylation using Phos-tag gels, which can resolve multiple phospho-isoforms of a target protein. Notably, apilimod treatment caused a change in the phosphorylation pattern of TFEB when separated by a Phos-tag gel that was comparable to torin1-treated cells (Fig. 2C). Importantly, lysates from *tfeb*^−/-^ RAW cells lacked most bands separated by Phos-tag gels and detected with anti-TFEB antibodies, identifying the signal as specific to TFEB (Fig. 2C). In addition, pre-incubation of lysates with λ phosphatase, collapsed all most bands signal into a single fast-running TFEB band (Fig. 2C), suggesting that we specifically detected a variety of phospho-TFEB isoforms. Overall, these data suggest that PIKfyve regulates kinases or phosphatases that impinge on TFEB, but these are seemingly in parallel and/or independent of mTOR.

### PIKfyve inhibition boosts lysosome gene expression but not lysosomal protein levels

Our observations suggest that PIKfyve may stimulate lysosomal gene and protein expression by activating TFEB and related factors. To test this, we measured mRNA levels of select lysosomal genes including LAMP1, MCOLN1, cathepsin D, and the V-ATPase H and D subunits using qRT-PCR. As expected, we observed augmented mRNA levels for these genes after 3 h incubation with apilimod (Fig. 3A). We next examined if the enhanced mRNA levels corresponded to an increase in protein levels. Surprisingly, the levels of the V-ATPase H subunit, LAMP1 and cathepsin D in apilimod-treated RAW macrophages were similar to control cells after 3 or 6 h of PIKfyve inhibition, though the small increase in LAMP1 protein levels at 6 h apilimod was statistically higher than resting cells (Fig. 3B, C). The decoupling between enhanced mRNA levels and unchanged protein levels may be due to the role of PIKfyve in targeting lysosomal proteins to lysosomes (Ikonomov et al., 2009a; Rutherford et al., 2006; Min et al., 2014). Given this, we suspected that TFEB activation may not contribute to lysosome swelling in cells acutely suppressed for PIKfyve.

**Figure 3:**
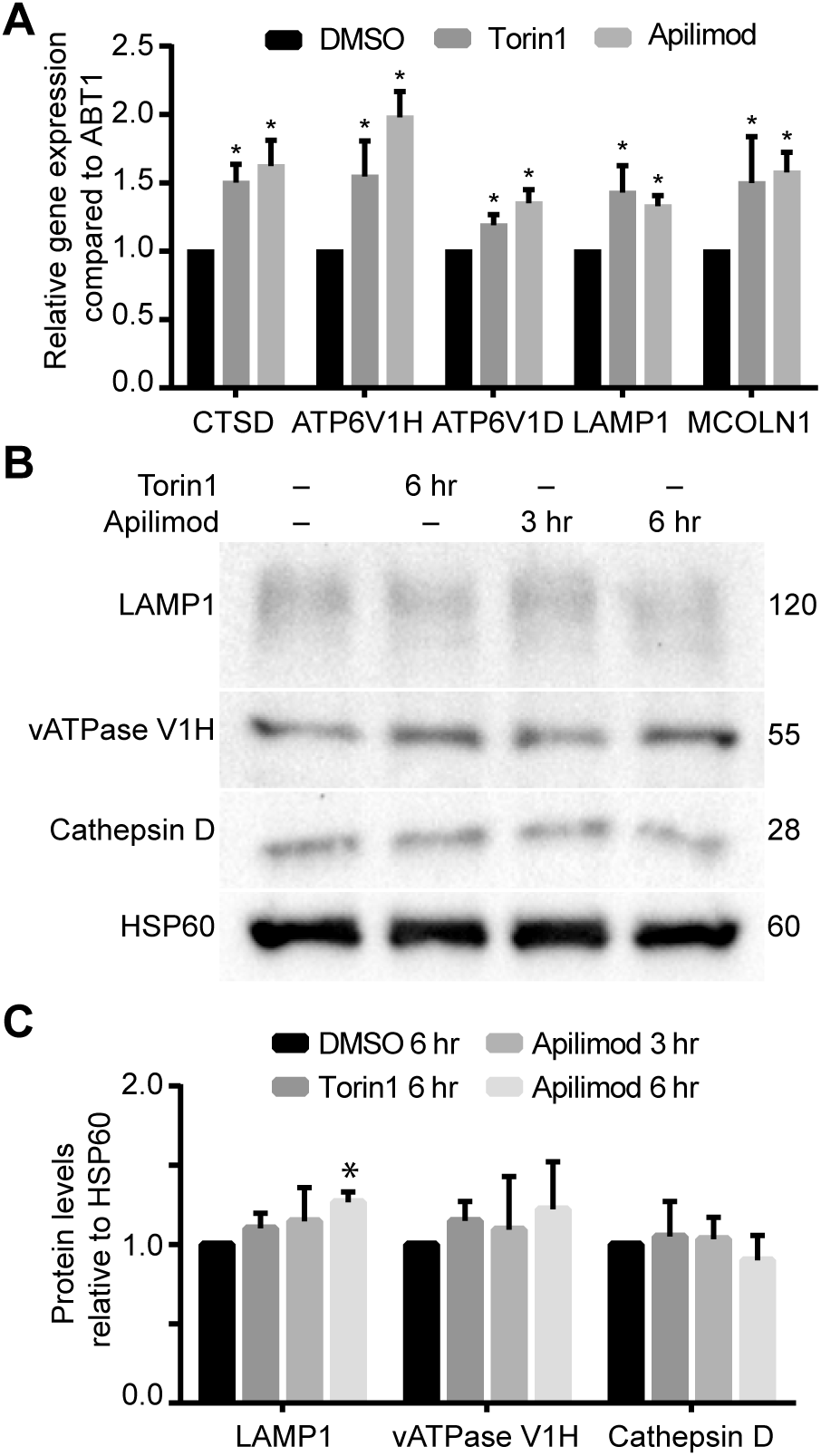
Acute PIKfyve inhibition enhances lysosomal gene transcription but not protein levels. A. Expression of select lysosomal genes quantified by qRT-PCR and normalized against Abt1. Shown is the mean ± SEM from seven independent experiments. B. Western blot against selected lysosomal proteins in RAW cells treated as indicated. Numbers on the right represent relative molecular weight (kDa). C. Quantification of indicated proteins demonstrated in B and shown as the mean intensity ± SEM from four independent blots. Asterisks (*) indicate statistical difference from respective control conditions (p<0.05) using multiple Student’s t-test with the Bonferroni correction.

### PIKfyve-inhibition induces lysosome swelling cells deleted for TFEB and related proteins

Next, we tested whether expression of TFEB and/or TFE3 is dispensable for lysosome swelling during PIKfyve inhibition. To do this, RAW macrophages deleted for TFEB, TFE3 or both (Pastore et al., 2016) were treated with vehicle or apilimod for 1 h and manually assessed for the average diameter and number of swollen vacuoles using differential interference contrast (DIC) optics. As suspected, RAW cells carrying deletions for TFEB and/or TFE3 were not impaired in apilimod-induced vacuolation, with vacuole number and size comparable to wild-type RAW cells (Supplemental Fig. S2A).

We then followed manual assessment with a more robust volumetric analysis to quantify the size and number of endo/lysosomes by decorating their luminal volume with fluid-phase pinocytic tracers like fluorescent-dextrans or Lucifer yellow. Notably, our labelling method likely decorates a collection of late endosomes, lysosomes and endolysosomes, a term that describes hybrid endosome-lysosome organelles (Huotari and Helenius, 2011). Thus, we use the term endo/lysosome to reflect the heterogeneous mixture of endosomes and lysosomes likely labelled here and to differentiate from endolysosomes. First, we showed that fluorescent-dextran co-localized to LAMP1-positive structures similarly well between all RAW strains, confirming that deletion of TFEB and/or TFE3 did not disrupt trafficking of the label to endo/lysosomes (Supplemental Fig. S2B). Second, we defined parameters such as thresholding and minimum particle size to estimate endo/lysosome number and volume (see Methods and Supplemental information for optimization testing; Supplemental Fig. S3). Using this approach, we conclude that the average number and volume of individual endo/lysosomes were similar between apilimod-treated wild-type and *tfeb*^*−/−*^, *tfe3*^*−/−*^ and *tfeb*^*−/−*^ *tfe3*^*−/−*^ RAW macrophages (Fig. 4A), consistent with the manual analysis (Supplemental Fig. S2A). Nevertheless, *tfeb*^*−/−*^ *tfe3*^*−/−*^ RAW macrophages retain expression of the related MITF protein, which becomes nuclear upon addition of apilimod (Supplemental Fig. S1); this raised the possibility that MITF may be sufficiently redundant with TFEB and TFE3 to promote lysosome expansion during PIKfyve inhibition. To test this, we opted to compare HeLa cells to counterpart HeLa cells deleted for all three proteins, previously described in (Nezich et al., 2015). Despite deletion of genes encoding TFEB, TFE3 and MITF, endo/lysosome enlargement caused by apilimod was indistinguishable between wild-type and mutant HeLa strains (Supplemental Fig. S4). Finally, we then asked whether protein biosynthesis was necessary for lysosomes to enlarge during PIKfyve repression. Strikingly, cycloheximide, which arrests protein biosynthesis, did not interfere with apilimod-induced lysosome swelling (Fig. 4B). As a control for the effectiveness of cycloheximide inhibition of protein synthesis, we assayed for translation activity with puromycylation and steady-state levels of p53, a high turnover protein. As shown in Fig. 4C, resting cells exhibited high-rate of puromycylation, which detects ribosome-mediated protein synthesis, whereas cycloheximide ablated this. In addition, cycloheximide rapidly depleted p53 levels, demonstrating that protein synthesis was arrested (Fig. 4D). Overall, our data suggest that protein biosynthesis and lysosome biogenesis controlled by TFEB and related transcription factors do not contribute substantially to endo/lysosome swelling during acute PIKfyve inhibition.

**Figure 4:**
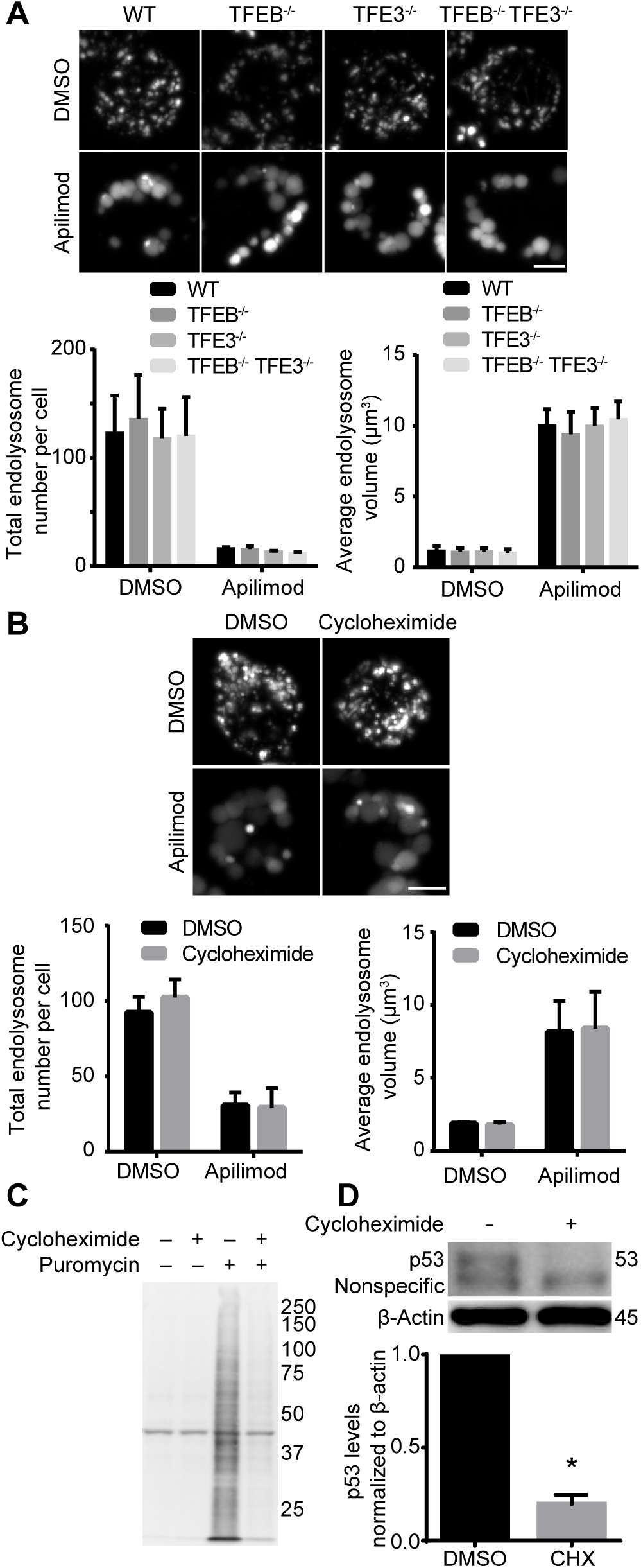
Volumetric analysis of endo/lysosomes in PIKfyve-curtailed wild-type, *tfeb*^−/-^, *tfe3*^−/-^ and *tfeb*^−/-^ *tfe3*^−/-^ RAW cells. A. Cells were pre-labelled with Lucifer yellow and then subjected to apilimod to induce lysosome vacuolation. Lysosome number and lysosome volume were quantified from 100 cells from three independent experiments. B. Lysosome morphology in wild-type RAW cells treated as indicated with cycloheximide. Lysosome volume and number were quantified. In all cases, data shown is the mean ±SEM from three independent experiments, with at least 100 cells per condition per experiment. C. Puromycylation of proteins in the absence and presence of cycloheximide. Puromycin is incorporated into elongating proteins by the ribosome, which can be detected by anti-puromycin antibodies during Western blotting. Cycloheximide-treatement obliterated puromycylation of proteins by blocking protein synthesis. The first two lanes resolved lysates unexposed to puromycin and thus accounts for non-specific bands detected by anti-puromycin antibodies. Representative blot shown from 3 independent experiments. D. Cycloheximide reduces p53 abundance relative to β-actin. Representative blot shown from 3 independent experiments. Numbers of the right represent relative molecular weight (in kDa). Data shown is the mean abundance ±SEM by measuring p53 signal as a ratio against β-actin. Asterisks (*) indicate statistical difference from respective control conditions (p<0.05) using Student’s t-test. Scale bar = 5 µm.

### PIKfyve inhibition concurrently increases endo/lysosome volume while decreasing endo/lysosome number

To better understand the process of endo/lysosome enlargement during PIKfyve inhibition, we further employed quantitative volumetric image analysis. Using this approach, we observed that the average volume of individual endo/lysosomes in RAW cells treated with apilimod increased with incubation period and relative to vehicle-only cells (Fig. 5A, B). Remarkably, the average number of endo/lysosomes per cell in apilimod-treated cells decreased considerably with incubation period and relative to vehicle-exposed cells (Fig. 5A, C). Despite the decrease in lysosome number, the total lysosome volume per cell remained unperturbed (Fig. 5A, D). As suggested above, we also observed similar trends with HeLa cells exposed to apilimod and *fig4*^*−/−*^ mouse embryonic fibroblasts; there was an increase in the average volume of individual endo/lysosomes, a concurrent decline in their number, while total lysosome volume per cell was not significantly altered (Supplemental Fig. S4, Fig. 5D-H). Finally, and as implied above, there was no significant difference in the average size, number and total volume of endo/lysosomes between wild-type RAW and HeLa cells and counterpart strains deleted for TFEB, TFE3 and/or MITF before and after PIKfyve suppression (Fig. 4; Supplemental Fig. S4). Interestingly, replacing apilimod-containing medium with fresh media reversed the effects on endo/lysosome volume and number per cell (Fig. 6). Overall, our results collectively suggest that acute PIKfyve inhibition not only enlarges endo/lysosomes, but also decreases their numbers without disturbing the total endo/lysosome volume.

**Figure 5:**
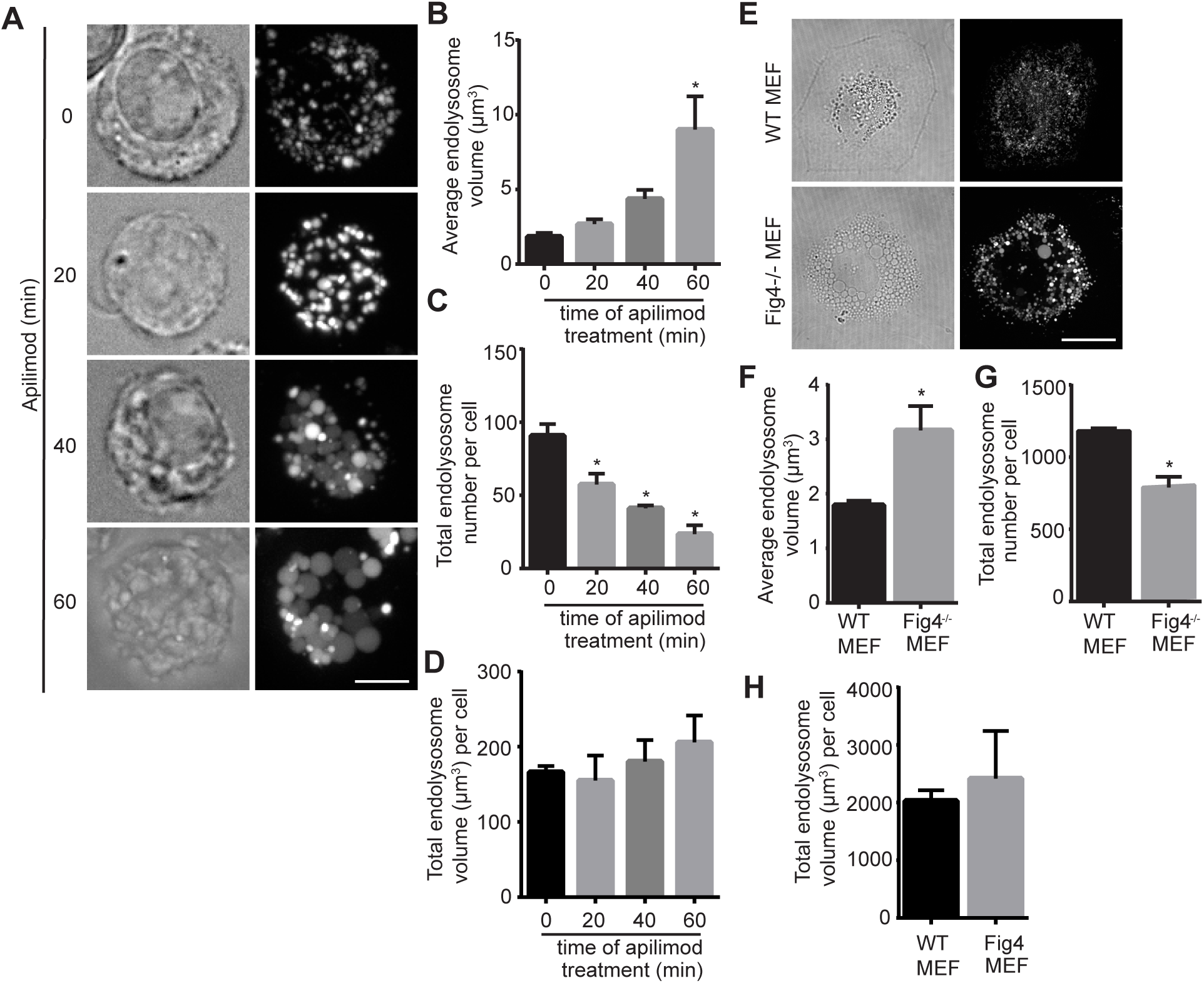
Volumetric analysis of endo/lysosomes during acute and chronic PIKfyve suppression. A. RAW cells were pre-labelled with Lucifer yellow as before and then exposed to vehicle-only or to 20 nM apilimod for the indicated times. Fluorescence micrographs are z-projections of 45-55 z-planes acquired by spinning disc confocal microscopy. Scale bar = 5 µm. B, C, D: Quantification of individual endo/lysosome volume (B), endo/lysosome number per RAW macrophage (C) and total endo/lysosome volume (D). E. Wild-type and *fig4*^*−/−*^ mouse embryonic fibroblasts labelled with Lucifer yellow. Scale bar = 30 µm. F,G, H. Analysis of individual endo/lysosome volume (F), endo/lysosome number per cell (G) and total endo/lysosome volume per cell (H) in wild-type and *fig4*^*−/−*^ mouse embryonic fibroblasts. In all cases, data shown are the mean ±SEM from three independent experiments, with at least 15-20 cells per condition per experiment. Asterisks (*) indicate statistical difference from respective control conditions (p<0.05) using one-way ANOVA and Tukey’s post-hoc test for data shown in B-D, and unpaired Student’s t-test for data displayed in E-H.

**Figure 6:**
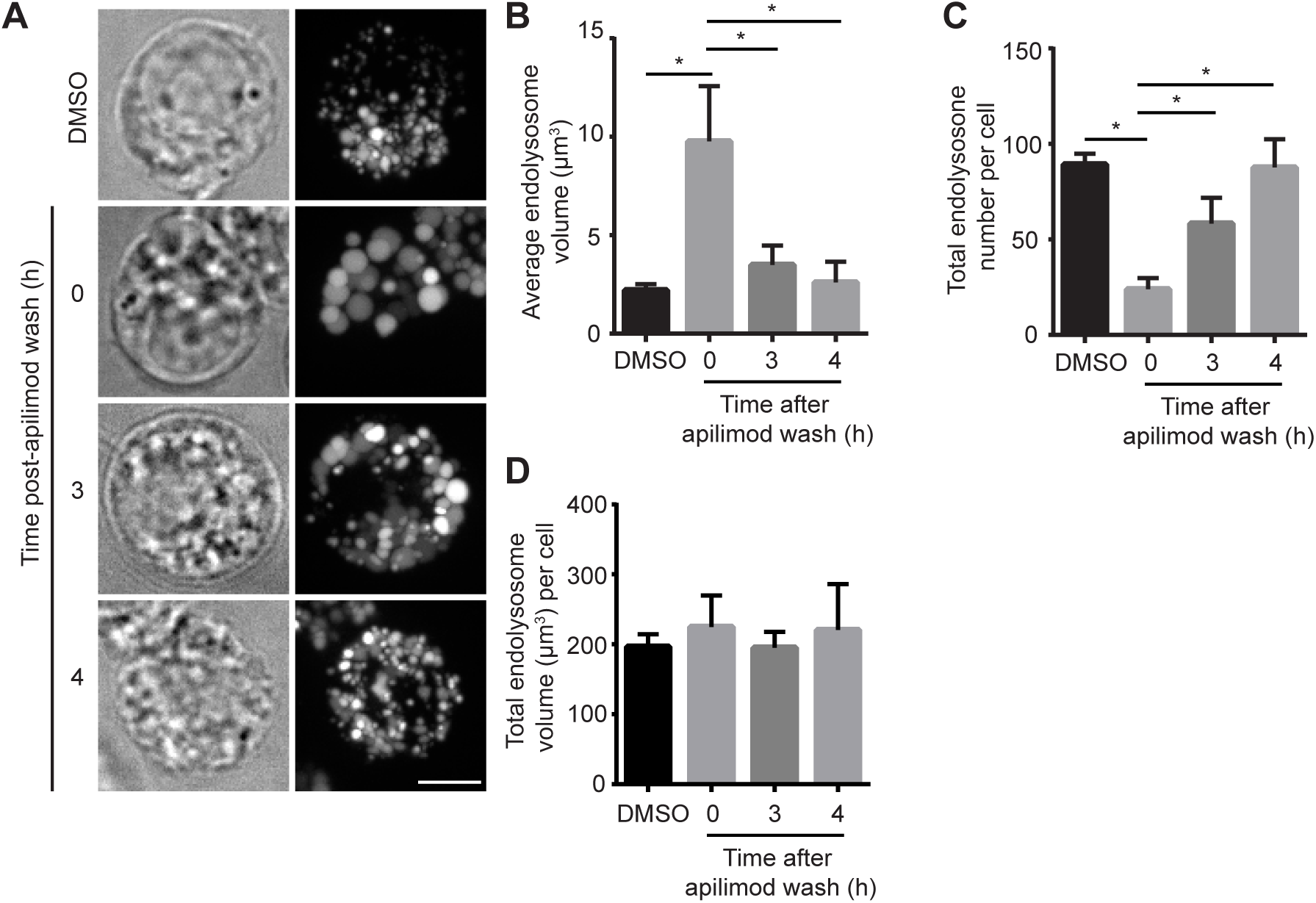
Volumetric analysis of endo/lysosome number and volume during PIKfyve reactivation. A. Cells were pre-labelled with Lucifer yellow as before and then exposed to vehicle-only or to 20 nM apilimod for 1 h, followed by removal and chase in fresh media for the indicated times. Fluorescence micrographs are z-projections of 45-55 z-planes acquired by spinning disc confocal microscopy. Scale bar = 5 µm. B, C, D: Volumetric quantification of individual endo/lysosome volume (B), endo/lysosome number per cell (C) and total endo/lysosome volume per cells (D). Data shown are the mean ±SEM from three independent experiments, with at least 15-20 cells per condition per experiment. Asterisks (*) indicates statistical difference between data indicated by lines (p<0.05) using one-way ANOVA and Tukey’s post-hoc test.

### Imbalanced fusion-fission cycles may underpin endo/lysosome swelling in PIKfyve-hindered cells

It is unlikely that endo/lysosome number is reduced during PIKfyve inhibition via lysosome lysis because the total lysosome volume per cell was unchanged relative to control cells. Instead, the reduction in endo/lysosome number coupled to their enlargement suggests that there is a shift towards endo/lysosome homotypic fusion relative to fission in PIKfyve-abrogated cells. Hence, we predicted that conditions that abate homotypic lysosome fusion would lessen enlargement and maintain higher lysosome numbers in PIKfyve-inhibited cells.

To test this hypothesis, we disrupted the microtubule system and motor activity using nocadozole and ciliobrevin, respectively, to impair lysosome-lysosome fusion (Rosa-Ferreira and Munro, 2011; Deng and Storrie, 1988; Jordens et al., 2001). Indeed, cells treated with these compounds resisted apilimod-induced swelling and retained higher endo/lysosome number relative to apilimod-only condition (Fig. 7A-C for nocadozole, Supplemental Fig. S5A-C for ciliobrevin). Conversely, endo/lysosome shrinkage was accelerated upon washing apilimod in microtubule-disrupted cells (Fig. 7D-F). To complement the pharmacological approach, we transfected cells with the p50 dynamitin subunit or dominant negative kinesin-1 to respectively impair dynein and kinesin. We observed reduced swelling and increased endo/lysosome numbers during apilimod exposure relative to untransfected cells (Supplemental Fig. S5D-I). Overall these data suggest that endo/lysosomes swell in PIKfyve-inhibited cells through endo/lysosome coalescence.

**Figure 7:**
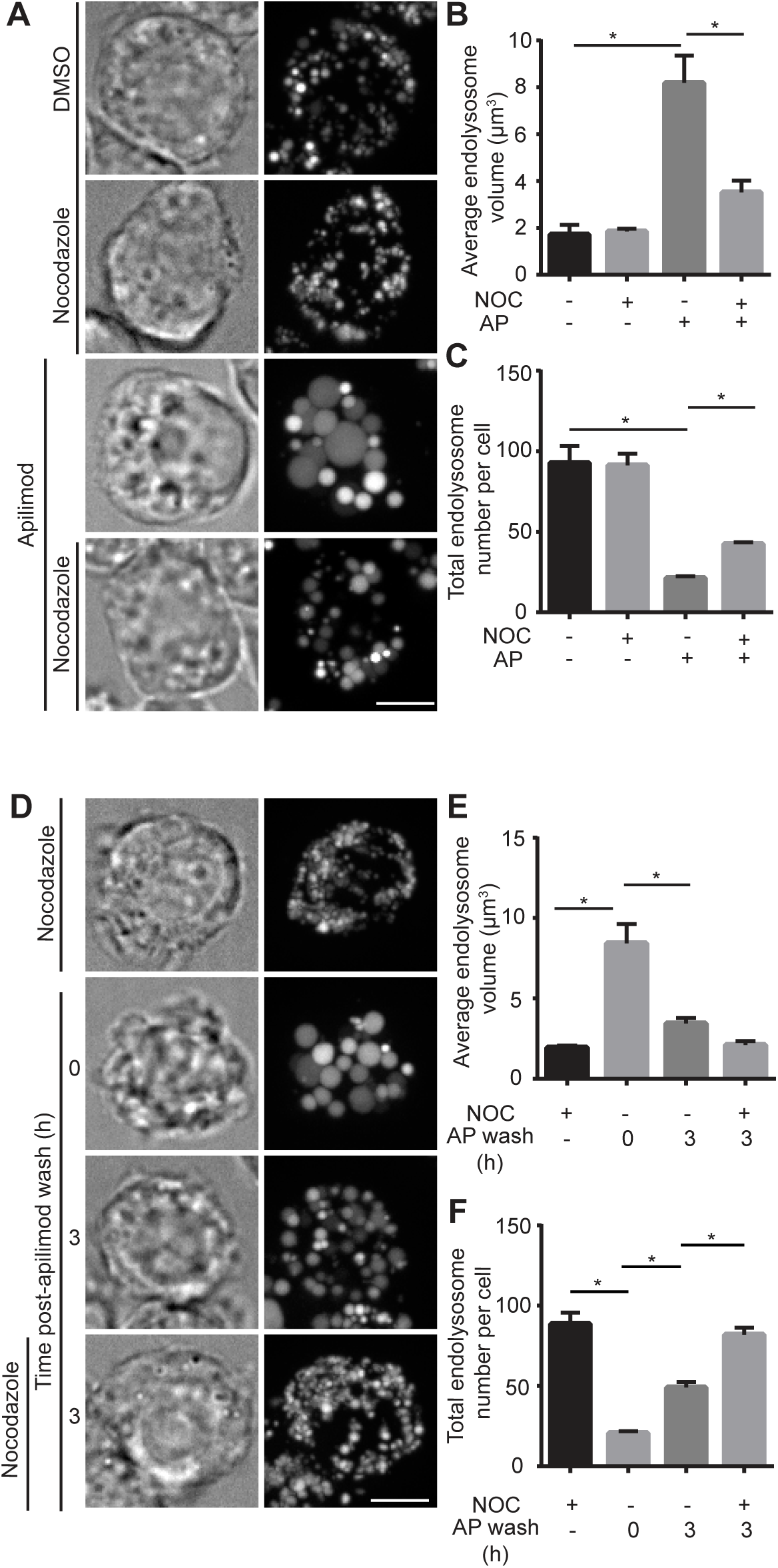
Volumetric analysis of endo/lysosome number and volume in cells disrupted for microtubule function. A. Cells were pre-labelled with Alexa^488^-conjugated dextran, followed by vehicle-alone, 10 µM nocadozole for 60 min, or co-treated with apilimod and vehicle or nocadozole for 1 h. Fluorescence micrographs are z-projections of 45-55 z-planes acquired by spinning disc confocal microscopy. B, C: Quantification of individual endo/lysosome volume (B) and endo/lysosome number per RAW macrophage (C). D. Cells were pre-labelled with Alexa^488^-conjugated dextran as before, followed by 20 nM apilimod for 1 h and then replaced with fresh media or media containing nocadozole for the indicated time period. E, F: Volumetric quantification of individual endo/lysosome volume (E) and endo/lysosome number per cell (D). In all cases, data shown are the mean ±SEM from three independent experiments, with at least 15-20 cells per condition per experiment. Asterisks (*) indicates statistical difference between conditions indicated by lines (p<0.05) using one-way ANOVA and Tukey’s post-hoc test. Scale bar = 5 µm.

Lysosomes undergo constant rounds of fusion and fission, including by “kiss-and-run” (Duclos et al., 2003; Bright et al., 2005; Pryor et al., 2000). We propose a model in which PIKfyve, likely through synthesis of PtdIns(3,5)P_2_, controls endo/lysosome fission. Consequently, in PIKfyve inhibited cells, lysosomes fail to fission and/or “kiss-and-marry”, resulting in endo/lysosome coalescence, enlargement, and a drop in total lysosome number. To support this model, we tried to visualize endo/lysosome swelling during apilimod exposure of RAW cells by using live-cell spinning disc confocal microscopy to obtain z-stacks at high temporal resolution. However, endo/lysosomes under observation failed to swell when acquiring z-stacks even as infrequently as every two minutes (Supplemental Fig. S6, Supplemental Movies 1 and 2). In addition, lysosomes appeared to become less dynamic as well. Indeed, when we moved to a different field-of-view within the same coverslip after the period of observation, endo/lysosomes were engorged (Supplemental Fig. S6). This suggests that photo-toxicity impairs apilimod-dependent endo/lysosome swelling. We could observe dynamic endo/lysosome movement and enlargement when imaging single-planes at 1 frame/2 min (Fig. 8A, B, Supplemental Movies 3 and 4). Unfortunately, this made tracking of individual endo/lysosomes difficult due to their highly dynamic nature.

**Figure 8:**
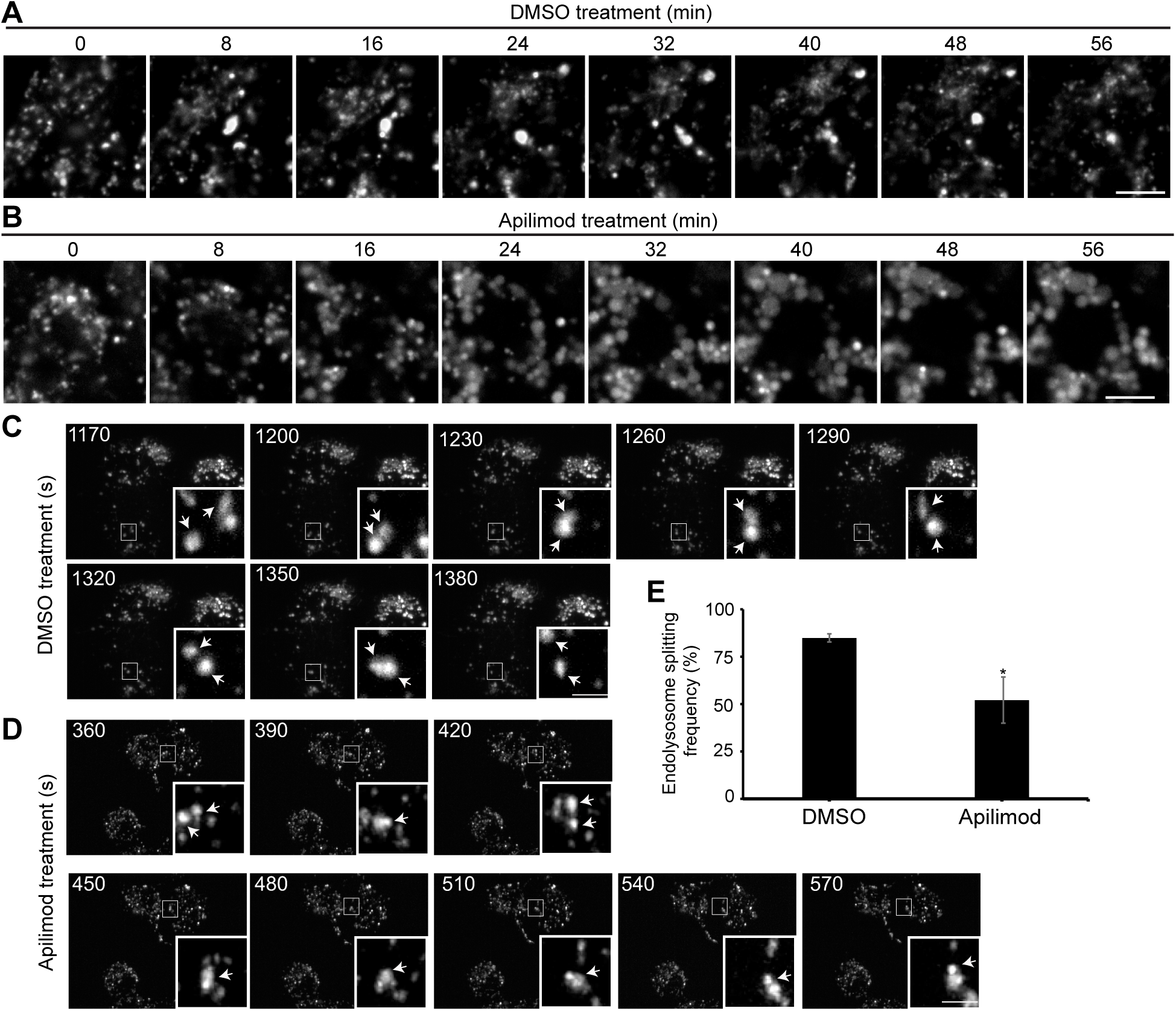
Live-cell imaging of apilimod-induced lysosome swelling. Cells were pre-labelled with Alexa^546^-conjugated dextran, followed by (A) vehicle-alone or (B) 20 nM apilimod at time 0 min. Single-plane imaging was effected by spinning disc microscopy at 1 frame/2 min. Time in min refers to time after adding vehicle or apilimod. See supplemental Movies 3 and 4 for full capture. Scale bar of image stills for (A) and (B) = 5 µm. Sweptfield imaging of RAW cells exposed to (C) vehicle or (D) 20nM apilimod. See supplemental Movies 5 and 6 for full capture. Shown are z-collapsed frames at 30 s intervals. The inset is a magnified portion of the field-of-view tracking a few endo/lysosomes (arrows) undergoing contact and split events, which may represent fusion or fission. Scale bar of image stills for (C) and (D) = 3 µm. E. Rate of “splitting” events over 30 min of vehicle or apilimod-treated cells showing a reduction in splitting in PIKfyve-inhibited cells, which may reflect reduced fission. Data were statistically analysed by Student’s t-test. Asterisk represents p<0.05.

To bypass the photo-toxicity issue, we switched to Sweptfield confocal microscopy, which employs dramatically less light energy. Using this technology, we could reconstruct cells at high-temporal resolution and using z-stacks. Indeed, we could track the dynamics of endo/lysosomes in control cells and those undergoing vacuolation in PIKfyve-inhibited cells over 2 h (Fig. 8C, D, Supplemental Movie S5 and S6). When imaging untreated cells, endo/lysosomes proved to be highly dynamic structures undertaking significant number of collisions, splitting and “tubulation” events (Fig. 8C, Supplemental Movie S5). After adding apilimod, endo/lysosomes underwent enlargement and became fewer, through coalescence (Fig. 8D, Supplemental Movie 6). However, it is possible to see that even enlarged endo/lysosome vacuoles in PIKfyve-suppressed cells retain their dynamic nature. To try to quantify fission and “run” events, we used particle tracking and splitting functions in Imaris as a proxy. Though this analysis cannot distinguish between fission and separation of overlapping but independent particles, we reasoned that this could estimate differences in fission rates if the defect is sufficiently large. Indeed, while approximately 85% of endo/lysosomes “split” in control cells over a period of 30 min, this was reduced to about 50% in apilimod-treated cells (Fig. 8E). Altogether, we propose that PIKfyve loss-of-function causes lysosome enlargement by reducing the rate of lysosome fission/”run” events relative to lysosome fusion/”kiss” events, leading to lysosome coalescence.

## Discussion

Disruption of PIKfyve activity causes several major physiological defects including neurological decay, lethal embryonic development, and inflammatory disease, among others (Ikonomov et al., 2011; Min et al., 2014; Chow et al., 2007; McCartney et al., 2014). It is unclear how these defects arise, but they likely relate to disruption of endo/lysosome homeostasis, including impaired hydrolytic function, reduced rates of autophagic flux, and endo/lysosome swelling (Martin et al., 2013; de Lartigue et al., 2009; Min et al., 2014; Kim et al., 2016). Endo/lysosome vacuolation remains the most dramatic and visible defect in PIKfyve-abrogated cells. Yet, the mechanism by which endo/lysosomes become engorged remains poorly defined. The prevailing model is that recycling from endo/lysosomes is impaired in PIKfyve/Fab1-defective cells, thus causing endo/lysosome and vacuole swelling (McCartney et al., 2014; Ho et al., 2012). Arguably, this is assumed to proceed through tubulo-vesicular transport intermediates formed during membrane fission catalysed by coat proteins like the retromer (Bryant et al., 1998; de Lartigue et al., 2009; Rutherford et al., 2006; Zhang et al., 2007). Here, we explored two other possibilities that could contribute to endo/lysosome enlargement: enhanced endo/lysosome biosynthesis and/or an imbalance in endo/lysosome fusion-fission dynamics. Collectively, our work intimates that enhanced lysosome biosynthesis does not drive endo/lysosome swelling. Instead, our work suggests that endo/lysosomes coalescence during PIKfyve inhibition, resulting in concurrent enlargement of endo/lysosomes and abatement in their numbers. We provide evidence that coalescence is due to a shift away from fission during endo/lysosome fusion-fission cycling (Supplemental Fig. S7). Our observations are consistent with recent work by Bissig *et al.* that intimate a role for PIKfyve in reforming terminal lysosomes from endolysosomes, a term they use to define a subset of lysosomes that are acidic and proteolytic (Bissig et al., 2017).

### PIKfyve and TFEB function

Inhibition of PIKfyve caused rapid and robust nuclear translocation of TFEB, TFE3 and MITF. Indeed, others have recently observed that TFEB shuttles into the nucleus in PIKfyve-inhibited cells (Gayle et al., 2017; Wang et al., 2015). Consistent with nuclear translocation, we also observed increased mRNA levels of selected lysosomal genes including LAMP1, V-ATPase subunits and cathepsins. This increase is in line with a recent study that looked at transcriptome-wide changes in mRNA levels in B-cell non-Hodgkins lymphoma cells treated with apilimod (Gayle et al., 2017). This suggests that PIKfyve regulates a negative-feedback loop to control lysosome biogenesis through suppression of TFEB and related factors. However, acute inhibition of PIKfyve led to either no change, or in the case of LAMP1, to a small increase in protein levels after 6 h of PIKfyve inhibition, despite the augmented mRNA levels. The decoupling between enhanced transcription and constant protein levels may result from pleiotropic effects of PIKfyve inhibition. For example, PIKfyve is known to govern trafficking of newly synthesized lysosomal genes by controlling recycling of mannose-6-phosphate receptors (Gayle et al., 2017; Ikonomov et al., 2009a; Rutherford et al., 2006; Zhang et al., 2007). By contrast, cells chronically impaired for PIKfyve have augmented LAMP1 levels (Ferguson et al., 2009, 2010). However, this LAMP1 accumulation is thought to occur through impaired autophagosome turnover rather than an increase in biosynthesis of LAMP1 (Ferguson et al., 2009; Kim et al., 2016). This may be consistent with the small increase in LAMP1 after prolonged inhibition of PIKfyve (Fig. 3C). Overall, given that endo/lysosome enlargement was unaffected in cells deleted for TFEB, TFE3 and/or MITF or treated with cycloheximide, we argue that biosynthesis does not significantly contribute to lysosome enlargement in cells acutely inhibited for PIKfyve, though we do not exclude a role in other PIKfyve-mediated functions.

### PIKfyve and TFEB regulation

TFEB and TFE3 are maintained in the cytosol via phosphorylation on specific residues that form docking sites for the 14-3-3 scaffolding protein (Roczniak-Ferguson et al., 2012; Martina et al., 2012). We now know that TFEB and related factors are subjected to multiple kinases including mTORC1, PKC, PKD, GSK3β and phosphatases such as the Ca^2+^-dependent calcineurin (Li et al., 2016b; Martina et al., 2014; Roczniak-Ferguson et al., 2012; Settembre et al., 2012; Najibi et al., 2016; Ploper et al., 2015; Marchand et al., 2015; Wang et al., 2015). Yet, mTORC1 regulation of TFEB remains as the best-understood regulatory circuit. Given that PIKfyve is needed for mTORC1 activity in adipocytes (Bridges et al., 2012), we examined the existence of a PIKfyve-mTORC1-TFEB axis. However, S6K remained phosphorylated in apilimod-treated cells, suggesting that mTORC1 was active. These observations are consistent with Wang *et al*. and Krishna *et al*. who also showed that mTORC1 retains its activity in PIKfyve-abrogated cells (Wang et al., 2015; Krishna et al., 2016). We considered the possibility that mTORC1 and starvation controls PIKfyve activity instead, which may then modulate TFEB, perhaps by a PtdIns(3,5)P_2_-dependent localization of TFEB to lysosomes, as has been shown for Tup1, a transcription factor in yeast that controls galactose metabolism (Han and Emr, 2011). Yet, we could not observe a significant decrease in PtdIns(3,5)P_2_ levels in mTOR-inhibited cells, though we cannot exclude disruption of a specific pool of PtdIns(3,5)P_2_ or of PtdIns(5)P, a lipid that is directly or indirectly synthesized by PIKfyve (Zolov et al., 2012; Sbrissa et al., 2012). Overall, our data suggest that mTORC1 and PIKfyve regulate TFEB in parallel. We speculate that PIKfyve may regulate other modulators of TFEB such as PKD, PKC and GSK3β.

### Enlargement of lysosomes in PIKfyve suppressed cells

Loss of PIKfyve function causes massive lysosome swelling. This has been partially explained through a defect in membrane recycling from endosomes and/or lysosomes (Ho et al., 2012; McCartney et al., 2014). This model arose through experiments done in yeast expressing an engineered ALP protein carrying a Golgi retrieval sequence (RS-ALP). RS-ALP traffics to the vacuole, where it is subjected to proteolytic maturation. Mature RS-ALP appears in the Golgi membrane fraction in wild-type yeast cells, but not in those defective in PtdIns(3,5)P_2_, suggesting a defect in retrieval from the vacuole (Bryant et al., 1998; Dove et al., 2004). In addition, PIKfyve appears to regulate retrieval of mannose-6-phosphate receptors from endo/lysosomes in mammalian cells, which suggests a link between PIKfyve and coat proteins like retromer (Ikonomov et al., 2009b; Rutherford et al., 2006; Zhang et al., 2007). Hence, endo/lysosome enlargement may partly occur through an imbalance in membrane influx, caused by unabated anterograde membrane flow and hindered recycling due to defects in forming tubulo-vesicular intermediates (Supplemental Fig. S7). However, our observations also suggest that the number of endo/lysosomes in PIKfyve-suppressed cells is significantly reduced. The diminished endo/lysosome number is unlikely due to lysosome lysis since the total endo/lysosome volume was unchanged in PIKfyve-arrested cells. Instead, we propose that PIKfyve suppression causes an imbalance in lysosome fusion-fission cycling, leading to lysosome coalescence and enlargement (Supplemental Fig. S7).

Lysosomes constantly interact with each other through full fusion and fission, or through “kiss-and-run” events. During the “kiss” event, at least two lysosomes dock and form a transient pore to help exchange and homogenize their luminal content (Bright et al., 2005; Luzio et al., 2009; Duclos et al., 2000). However, complete fusion and coalescence is pre-empted by the “run” event, when the two lysosomes separate, thus preventing coalescence. Our work suggests that PIKfyve, likely through PtdIns(3,5)P_2_, plays a role in mediating lysosome fission to balance fusion. This role may occur during complete fusion-fission cycles and/or by catalysing the “run” event during “kiss-and’run” (Supplemental Fig. S7). In either case, failure to fission would result in overt coalescence, resulting in lysosome enlargement and reduced numbers. This model is consistent with recent work by Bissig *et al.* that intimate a role for PIKfyve in reforming terminal lysosomes from endolysosomes (Bissig et al., 2017). We do not understand how PIKfyve modulates lysosome fission dynamics. However, we speculate that mechanisms that offset lysosome coalescence are distinct from those that drive vesiculation and recycling (Krishna et al., 2016; Bryant et al., 1998; Rutherford et al., 2006). One interesting possibility may relate to contact sites between endo/lysosome and the endoplasmic reticulum since these have been shown to demarcate and assemble molecular machines that catalyse endolysosome fission (Friedman et al., 2013; Rowland et al., 2014). Thus, it is appealing to speculate that PIKfyve and PtdIns(3,5)P_2_ may mediate this process or be regulated by these contact sites to drive endo/lysosome fission.

Overall, we examined two possible mechanisms that might have contributed to endo/lysosome enlargement in PIKfyve-inhibited cells. While TFEB and related factors are controlled by PIKfyve, likely establishing a negative-feedback loop for lysosome biogenesis, we did not obtain evidence to suggest that biosynthesis contributes to lysosome swelling during acute PIKfyve inhibition. Instead, we observed that loss of PIKfyve activity caused concomitant reduction in the number of and increase in the volume of endo/lysosomes, suggesting that endo/lysosomes undergo coalescence through fusion and impaired fission. Future work should delineate the contribution of lysosome coalescence and vesicular recycling to lysosome enlargement.

## Materials and Methods

### Cell culture and transfection

RAW 264.7 macrophages strains and HeLa cell strains were cultured in Dulbecco’s Modified Eagle Medium (DMEM; Wisent, St. Bruno, QC) supplemented with 5% heat inactivated fetal bovine serum (FBS; Wisent). Mouse embryonic fibroblast (MEF) cells were cultured in Roswell Park Memorial Institute media (RPMI-1640: Gibco, Burlington, ON) with 15% FBS. All cells were authenticated, cleared of contamination and were cultured at 37°C with 5% CO_2_. Transient transfections were performed with FuGene HD (Promega, Madison, WI) following manufacturer’s instructions, using a 3:1 DNA to transfection reagent ratio. The transfection mixture was replaced with fresh media after 4-6 h and cells were employed 24 h post-transfection. Silencing RNA oligonucleotides were electroporated into RAW cells with the Neon Electroporation system (Life Technologies, Burlington, ON) following manufacturer’s instructions. Briefly, two rounds of knockdown were performed using ON-TARGETplus PIKfyve SMARTpool siRNA (L-040127-00-0005) or ON-TARGETplus non-targeting control pool (GE Healthcare, Mississauga, ON) over 48 h.

### Pharmacological manipulation of cells

Cells were treated with apilimod (Toronto Research Chemicals, Toronto, ON) at indicated concentrations and times. Torin-1 (Tocris Bioscience, Minneapolis, MN) or rapamycin (BioShop, Burlington, ON) were used as positive controls for TFEB nuclear localization and mTORC1 inhibition. Cells were also incubated with 100 µM ciliobrevin D (EMD Millipore, Toronto, ON) or 10 µM nocodazole (Sigma-Aldrich, Oakville, ON).

### Western blotting

Whole cell lysates were generated with 2x Laemmli sample buffer supplemented with protease inhibitor cocktail (Sigma-Aldrich) and PhosSTOP phosphatase inhibitor (Roche, Laval, QC) where necessary. Lysates were passed through a 27-gauge needle 6x and heated. Puromycylation assays were carried out with the addition of 10 µg/mL puromycin for 15 minutes before cell lysis. When protein dephosphorylation was required, cells were lysed with 1% triton-X100 in PBS supplemented with protease inhibitor cocktail, spun at 16,000 x *g* for 10 minutes, and the supernatant was subjected to lambda protein phosphatase (NEB, Whitby, ON) as per manufacturer’s instruction. Briefly, 20 µg of protein was incubated with 50 mM HEPES, 100 mM NaCl, 2 mM DTT, 0.01% Brij 25, 1 mM MnCl_2_, and 200 units of phosphatase at 30°C for 60 min. The reaction as stopped with 5x Laemmli sample buffer and heated.

Most samples were subjected to SDS-PAGE through a 10% acrylamide resolving gel. To assay for TFEB mobility shift, we used an 8% acrylamide resolving gel supplemented with 12.5 μM Phos-tag (Wako Chemicals USA, Richmond, VA) and 25 μM MnCl_2_. Phos-tag gels were washed three times with 100 mM EDTA for 10 min each, followed by a wash with Transfer buffer (25 mM Tris-HCl, 192 mM glycine, 20% methanol). Proteins were transferred to PVDF, blocked and incubated with primary and HRP-linked secondary antibodies using Tris-buffered saline plus 0.1% Tween-20 with either 5% skim milk or 3% BSA. Proteins were visualized using Clarity enhanced chemiluminescence (Bio-Rad Laboratories, Mississauga, ON) with a ChemiDoc XRS+ or ChemiDoc Touch imaging system (Bio-Rad). Quantification of proteins were determined using Image Lab software (Bio-Rad) by normalizing against a loading control and then normalizing against the vehicle-treated control. Rabbit monoclonal antibodies used were against cathepsin D (1:1000, GTX62603, GeneTex Inc., Irvine, CA), HSP60 (1:1000, 12165S, Cell Signaling Technologies, Danvers, MA), and p70 S6 kinase (1:1000, 2708S, Cell Signaling). Rabbit polyclonal antibodies were used against ATP6V1H (1:1000, GTX110778, GeneTex), TFEB (1:2000, A303-673A, Bethyl Laboratories, Montgomery, TX), phospho^Thr389^-p70 S6 kinase (1:1000, 9205S, Cell Signaling), and β-actin (1:1000, 4967S, Cell Signaling). Rat monoclonal antibodies were used against LAMP1 (1:200, 1D4B, Developmental Studies Hybridoma Bank, Iowa City, IO). Mouse monoclonal antibodies were used against puromycin (1:1000, 540411, Millipore, Burlington, MA), and p53 (1:1000, 2524S, Cell Signaling). Secondary HRP-linked antibodies were raised in donkey (Bethyl).

### Quantitative RT-PCR

Total RNA was extracted from RAW cells using the GeneJET RNA purification kit (Thermo Fisher Scientific, Mississauga, ON). Equal quantities of mRNA were reverse transcribed with SuperScript VILO cDNA synthesis kit (Invitrogen) following manufacturer’s guidelines. The subsequent cDNA was diluted 1:100 and amplified for quantitative PCR using the TaqMan Fast Advanced Master Mix (Applied Biosystems, Foster City, CA) in the presence of TaqMan assays with a Step One Plus Real-Time PCR thermal cycler (Applied Biosystems) with Step One software (version 2.2.2; Applied Biosystems). The TaqMan gene expression assays (Invitrogen) for the reference gene Abt1 (Mm00803824_m1) and target genes Ctsd (Mm00515586_m1), Atp6v1h (Mm00505548_m1), ATP6V1D (Mm00445832_m1), LAMP1 (Mm00495262_m1), and Mcoln1 (Mm00522550_m1) were done in triplicate. Target gene expression was determined by relative quantification (ΔΔCt method) to Abt1 and the vehicle-treated control sample.

### Endo/lysosomal labelling

Endo/lysosomes in RAW cells, HeLa cells and MEFs were labelled by incubating cells with either 200 µg/mL Alexa^488^- or Alexa^546^-conjugated dextran (ThermoFisher) or with 2.5 mg/mL Lucifer yellow (ThermoFisher) in cell-specific complete medium for 2 h at 37 ºC in 5% CO_2_, followed by washing in phosphate-buffered saline and replenishing with fresh complete medium for 1 h before live-cell imaging. This method likely labels a spectrum of late endosomes, lysosomes and endolysosomes, which are endosome-lysosome hybrids (Huotari and Helenius, 2011). Thus, we refer to dextran- or Lucifer yellow-labelled structures as endo/lysosomes.

### Epifluorescence and spinning disc confocal microscopy

Manual quantification of lysosome size and imaging of GFP-based transfections were imaged using a Leica DM5000B system outfitted with a DFC350FX camera and controlled by Leica application suite (Leica Microsystems Inc., Concord, ON). TFEB-GFP transfected into PIKfyve knockout cells and endogenous TFEB in RAW cells were imaged using an Olympus IX83 microscope with a Hamamatsu ORCA-Flash4.0 digital camera controlled by CellSens Dimensions software (Olympus Canada Inc., Richmond Hill, ON). Endogenous TFEB was further deconvolved using a constrained iterative in CellSens Dimensions. All other imaging of fixed cells was done with a Quorum DisKovery Spinning disc confocal microscope system equipped with a Leica DMi8 microscope connected to Andor Zyla 4.2 Megapixel sCMOS camera and controlled by Quorum Wave FX Powered by MetaMorph software (Quorum Technologies, Guelph, ON). Live-cell imaging by spinning disc confocal microscopy was accomplished with either an Olympus IX81 inverted microscope equipped with a Hamamatsu C9100-13 EMCCD camera and a 60X 1.35 N.A. objective and controlled with Volocity 6.3.0 (PerkinElmer, Bolton, ON). Alternatively, we employed the Quorum DisKovery Spinning disc confocal microscope system above but using the iXON 897 EMCCD camera. Cells were imaged live using cell-specific complete medium in a 5% CO_2_ chamber at 37×C. All microscopes were equipped with standard filters appropriate to fluorophores employed in this study.

### ^3^H-myo-inositol labelling of phosphoinositides and phosphoinositide measurement by HPLC-coupled flow scintillation

RAW cells were incubated for 24 h in inositol-free DMEM (MP Biomedical, CA) with 10 µCi/mL myo-[2-^3^H(N)] inositol (Perkin Elmer, MA), 10% dialyzed FBS (Gibco), 4 mM L glutamine (Sigma Aldrich), 1x insulin-transferrin-selenium-ethanolamine (Gibco), and 20 mM Hepes (Gibco). Cells were then treated with torin1 or apilimod for 1 h. Reactions were arrested in 600 µL of 4.5% perchloric acid (v/v) on ice for 15 min, scraped, and pelleted at 12,000 x *g* for 10 minutes. Pellets were washed with 1 mL of ice cold 0.1 M EDTA and resuspended in 50 µL of water. Phospholipid deacylation was then done in 500 µL of methanol/40% methylamine/1 butanol (45.7% methanol: 10.7% methylamine: 11.4% 1-butanol (v/v)) for 50 min at 53°C. Samples were vacuum-dried and washed twice by resuspending in 300 µL of water and drying. Dried samples were resuspended again in 450 µL of water and extracted with 300 µL of 1butanol/ethyl ether/ethyl formate (20:4:1), vortexed for 5 min and centrifugation at 12,000x g for 2 min. The bottom aqueous layer was collected and extracted two more times. The aqueous layer was dried in vacuum and resuspended in 50 µL of water. Equal counts of ^3^H were separated by HPLC (Agilent Technologies, Mississauga, ON) through an anion exchange 4.6 × 250-mm column (Phenomenex, Torrance, CA) with a flow rate of 1 mL/min and subjected to a gradient of water (buffer A) and 1 M (NH_4_)_2_HPO_4_, pH 3.8 (adjusted with phosphoric acid) (buffer B) as follows: 0% B for 5 min, 0 to 2% B for 15 minutes, 2% B for 80 minutes, 2 to 10% B for 20 minutes, 10% B for 30 minutes, 10 to 80% B for 10 minutes, 80% B for 5 minutes, 80 to 0% B for 5 minutes. The radiolabeled eluate was detected by β-RAM 4 (LabLogic, Brandon, FL) with a 1:2 ratio of eluate to scintillant (LabLogic) and analyzed by Laura 4 software. Phosphoinositides were normalized against the parent phosphatidylinositol peak.

### Live-cell imaging with swept-field confocal microscopy

Cells were seeded into single-glass bottom micro-well dishes (MatTek, Ashland, MA) and grown to 70%-80% confluency. Cells were then incubated for 2 hours in 200 µL DMEM medium supplemented with 10% FBS with a 200 µg/mL of Alexa^555^-conjugated dextran (ThermoFisher). Samples were washed and incubation media was replaced with fresh complete DMEM followed by a 90 min post-incubation period. Live cell imaging was performed with a Nikon Ti inverted microscope equipped with a Prairie Technologies swept-field slit scanning confocal scan head and a 100x 1.49 N.A. objective (Prairie Technologies, Sioux Falls, SD). Nikon NIS Elements software equipped with a Photometrics Prime 95B back illuminated sCMOS camera was used for image acquisition while high speed triggered acquisition was controlled by a National Instruments DAQ card and Mad City Labs Piezo Z stage controller. Sample volumes were acquired at 100 ms per z-slice at 30 s intervals over a 30 min observation period before adding 20 nM apilimod and then 2 h in the presence of apilimod.

### Immunofluorescence

Following treatments, cells were fixed with 4% paraformaldehyde (v/v) for 15 min. Cells were blocked and permeabilized with 0.2% (v/v) Triton-X and 2% (v/v) BSA for 10 min, then subjected to rabbit polyclonal antibody against TFEB (1:250; Bethyl) and Dylight-conjugated donkey polyclonal antibody against rabbit (1:500; Bethyl). Nuclei were counter stained with 0.4 μg/mL of DAPI. For LAMP1 specific immunostaining, cells were fixed and permeabilized with 100% ice cold methanol for 5 minutes, followed by a 2% BSA block, incubation with Dylight-conjugated rat monoclonal antibody against LAMP1 (1:200, Clone 1D4B; Developmental Studies Hybridoma Bank) and polyclonal donkey anti-rat IgG antibodies (1:500; Bethyl).

### Imaging analysis and volumetrics

To quantify the proportion of cells with nuclear TFEB-GFP (including TFE3-GFP or MiTF-GFP), RAW cells were visually scored as cytosolic if the cytosol had equal or greater intensity than the nucleus. Alternatively, the nuclear:cytosolic ratio of endogenous TFEB was measured as the ratio of the mean intensity in the nucleus over the mean intensity in the cytosol using ImageJ. To manually quantify vacuole diameter in DIC images, vacuoles were first defined as having a diameter greater than 1.5 µm and then the diameter was measuring using line plot in ImageJ.

To quantify average endo/lysosome volume and number per cells, we employed volumetric and particle detection tools in Volocity (Volocity 6.3.0) or Icy BioImage Analysis. Z-stack images were imported into the software and subject to signal thresholding. First, we tried thresholding at 1.5, 2x, and 2.5x of average cytosolic fluorescence background. This was then subject to particle detection by defining particles as those with a volume > 0.3 µm^3^, which helped eliminate noise-derived particles. Regions of interest where then drawn around each cell for analysis and particles split using watershed function. Importantly, while increasing thresholding levels reduced absolute number of particles, particle size and total particle volume, i) the reduction occurred at similar levels between control and apilimod-treated cells and ii) the relative trends between control and apilimod-treated cells remained the same (Supplemental Fig. S3A, B). Hence, we employed a threshold of 2x the average cytosol fluorescence intensity for remaining analysis. Analysis of endo/lysosome splitting frequency was performed using Imaris (Bitplane, Concord, MA) using its *ImarisTrackLineage* module, where endo/lysosome splitting was defined as the frequency of events producing two particles from a single apparent particle. Image adjustments such as enhancing contrast was done with ImageJ or Adobe Photoshop (Adobe Systems, San Jose, CA) without altering the relative signals within images. Figures were assembled with Adobe Illustrator (Adobe Systems).

### Statistical analysis

All experiments were repeated at least three independent times. The respective figure legends indicate the total number of experiments and number of samples/cells assessed. The mean and measure of variation (standard deviation) or error (standard of error of the mean) are indicated. All experimental conditions were statistically analysed with either am unpaired Student’s t-test when comparing two parameters only or with a one-way ANOVA test when comparing multiple conditions in non-normalized controls. ANOVA tests were coupled to Tukey’s post-hoc test to analyse pairwise conditions. For multiple comparisons with normalized controls, Student’s t-tests were utilized with Bonferroni correction. Here, we accept p<0.05 as indication of statistical significance.

## Acknowledgements

Plasmid constructs expressing the kinesin-1 dominant negative and of the p50 dynamitin subunit were obtained from Addgene (Cambridge, MA). Plasmids encoding GFP fusion proteins of TFEB, TFE3 and MITF were kind gifts from Dr. Shawn Ferguson at Yale University. The *tfeb*^−/-^ *tfe3*^−/-^ *mitf*^−/-^ HeLa cells were a kind gift from Dr. Richard Youle at the National Institutes of Health.

## Authors contributions

CHC and GS contributed equally to this work, designing and performing experiments, analysing data and co-write the manuscript. MAG, RMD, ZAO performed and analysed experiments. CW and SCW performed imaging with sweptfield microscopy. GL extracted mouse embryonic fibroblasts. RP provided reagents and intellectual contributions to the manuscript. RJB acquired funding, designed experiments and co-wrote the manuscript.

## Abbreviations

DIC: Differential interference contrast
DMSO: Dimethyl sulfoxide
LY: Lucifer yellow
MEFs: mouse embryonic fibroblasts
PtdInsP: phosphoinositides
PtdIns(3)P: phosphatidylinositol-3-phosphate
PtdIns(3,5)P_2_: phosphatidylinositol-3,5-bisphosphate
STD: standard deviation
SEM: standard error of the mean

